# State-Transition Analysis of Time-Sequential microRNA Expression Predicts Development of Acute Myeloid Leukemia

**DOI:** 10.1101/2021.03.02.433582

**Authors:** D. E. Frankhouser, D. O’Meally, S. Branciamore, L. Uechi, H. Qin, X. Wu, N. Carlesso, G. Marcucci, R. C. Rockne, Ya-Huei Kuo

## Abstract

MicroRNAs (miRNAs) are small non-coding RNA molecules involved in post-transcriptional regulation of gene expression and have been shown to hold prognostic value in a variety of settings, including acute myeloid leukemia (AML). However, the temporal dynamics of miRNA expression profiles as it relates to AML initiation and progression is poorly understood. Using serial samples from a mouse model of AML, we show that the miRNA transcriptome undergoes state-transition during AML initiation and progression. The AML state-transition was visualized and modeled by constructing an AML state-space from singular value decomposition of the time-series miRNA sequencing data. Within the AML state-space, we identified critical points of AML development characterized by unique differentially expressed miRNAs compared to healthy controls at critical points of leukemogenesis (early, transition, and late). Interestingly, we observed that changes in miRNA expression during leukemogenesis followed two patterns: 1) a monotonic pattern with continuously increasing or decreasing expression; and 2) a non-monotonic pattern with a local maximum or minimum at the transition critical point which was the “point of no-return” from health to AML. We validated the AML state-space and dynamics in an independent cohort of mice and demonstrated the state-transition model accurately predicted time to AML. Of note, we show that the miRNA-derived state-transition model produced a state-space and critical points that were strikingly similar, but not identical to that produced by the coding (i.e., messenger [m]RNA-based) transcriptome. This indicates that while both miRNA and mRNA expression may provide similar information, they also capture independent features of AML state-transition.

**Significance:** We show that the microRNA transcriptome undergoes a global state transition during the initiation and progression of acute myeloid leukemia, and accurately predicts time to disease development.

## Introduction

Acute myeloid leukemia (AML) is a molecularly heterogeneous neoplastic disease originating in the bone marrow (BM) with more than 20,000 new cases diagnosed in the USA each year^1^. The relatively low 5-year survival rate of 28% reflects the urgent need for more effective treatments. With the rapid development and pervasive use of sequencing technologies in the clinical management of AML, there is an opportunity to take advantage of time-sequential multi-omic samplings of relevant tissues (BM and peripheral blood [PB]), identify new targets and devise novel therapeutic approaches.

State-transition models have been a useful way to interpret and predict time-sequential dynamics in stochastic biological systems. The biological applications of state-transition models include development, cell differentiation, and disease. In the context of diseases, including cancer, state-transitions are useful for studying disease initiation and progression. Constructing a state-space to model biological transitions can capture changes produced by a vast number of processes that occur simultaneously in a biological system. State-spaces have been constructed using a number of different types of data, but the mRNA transcriptome is often used because it is easy to assay and contains sufficient information to represent the main cellular processes. However, the large amount of information and high dimensionality of the transcriptome also presents challenges. Dimensionality reduction techniques, such as SVD used here and generalized SVD for multi-omic and pan-cancer studies, have proven useful because they represent the information of the entire system as linear combinations of orthogonal basis vectors^2–4^. Similar mathematical approaches including endogenous network theory have been used to identify steady states, or attractors, using configurations of gene regulatory networks^5–9^.

MicroRNAs (miRNA) are small non-coding RNA molecules involved in post-transcriptional regulation of gene expression. MiRNA expression profiles have been associated with pathogenesis and prognosis of AML. Yet very little is known about the dynamics of miRNA expression over the course of AML initiation and progression and how it can be targeted therapeutically. To our knowledge, state-transition modeling of miRNA transcriptome in AML as not been previously reported.

Here, using PB mononuclear cells (PBMCs) collected at sequential timepoints from a murine model of inv(16) AML, we show that miRNA transcriptome undergoes a state-transition (i.e., occupies stepwise transcriptional states) from disease initiation to progression. We defined a state-space and identified critical points which represent phenotypic states of AML progression. Analysis of the critical points identified miRNA-based regulatory events that predicts the dynamics of AML development from health to overt disease. This approach allowed us also to prioritize miRNAs that have a concerted role in AML development and that can potentially represent novel therapeutic targets.

## Materials and Methods

### Mouse model

The expression of the leukemogenic fusion gene *Cbfb-MYH11 (CM)* in conditional *CM* knock-in mice (*Cbfb*^*56M/+*^*/Mx1-Cre)* leads to development of AML with a median survival of approximately 4 months after induction of *CM*. This model recapitulates the human inv(16) AML, one of the common subsets of AML characterized by the rearrangement of chromosome 16 at bands p13 and q22, which at the molecular level creates the chimeric fusion gene *CBFB-MYH11*. To induce CM expression, 6-8 weeks old *CM* knock-in mice were injected intraperitoneally with polyinosinic–polycytidylic acid [poly (I:C)] (InvivoGen, tlrl-picw-250) at 14 mg/kg/dose every other day for a total of 7 doses. Age-matched littermates lacking the transgene were similarly treated and used as control. All mice were maintained in an Association for Assessment and Accreditation of Laboratory Animal Care–accredited animal facility and all experimental procedures were performed in accordance with federal and state government guidelines and established institutional guidelines and protocols approved by the Institutional Animal Care and Use Committee at City of Hope.

### Experimental design

We used two independent cohorts of mice as training and validation cohorts. For the training cohort, we collected PBMC from the *CM*-induced mice (n = 7) and the similarly treated littermate controls (n = 7) before induction (t = 0) and monthly after induction up to 10 months (t = 1-10) or when the mouse developed leukemia and became moribund, whichever event occurred first. Similarly, for the validation cohort, we collected PBMC from *CM* mice (n=9) or control (n=7) before induction and monthly thereafter up to 6 months. Total RNA was isolated from PBMC using AllPrep DNA/RNA Kit (Qiagen). Sequencing details including library preparation and differential expression analysis can be found in the supplemental methods.

### Mathematical model of state-transition

Using a state-transition model, we represented the miRNA transcriptome of each individual mouse as a particle in a “leukemogenic” double well quasi-potential with two steady states: normal hematopoiesis (health; *c*_1_) and AML (*c*_3_) separated by an unstable transition state (*c*_2_) (**Figure 1A**). After induction of the leukemogenic CM gene, we postulated that the potential energy landscape was perturbed such that a transition from health to leukemia became more likely.

**Figure 1.**
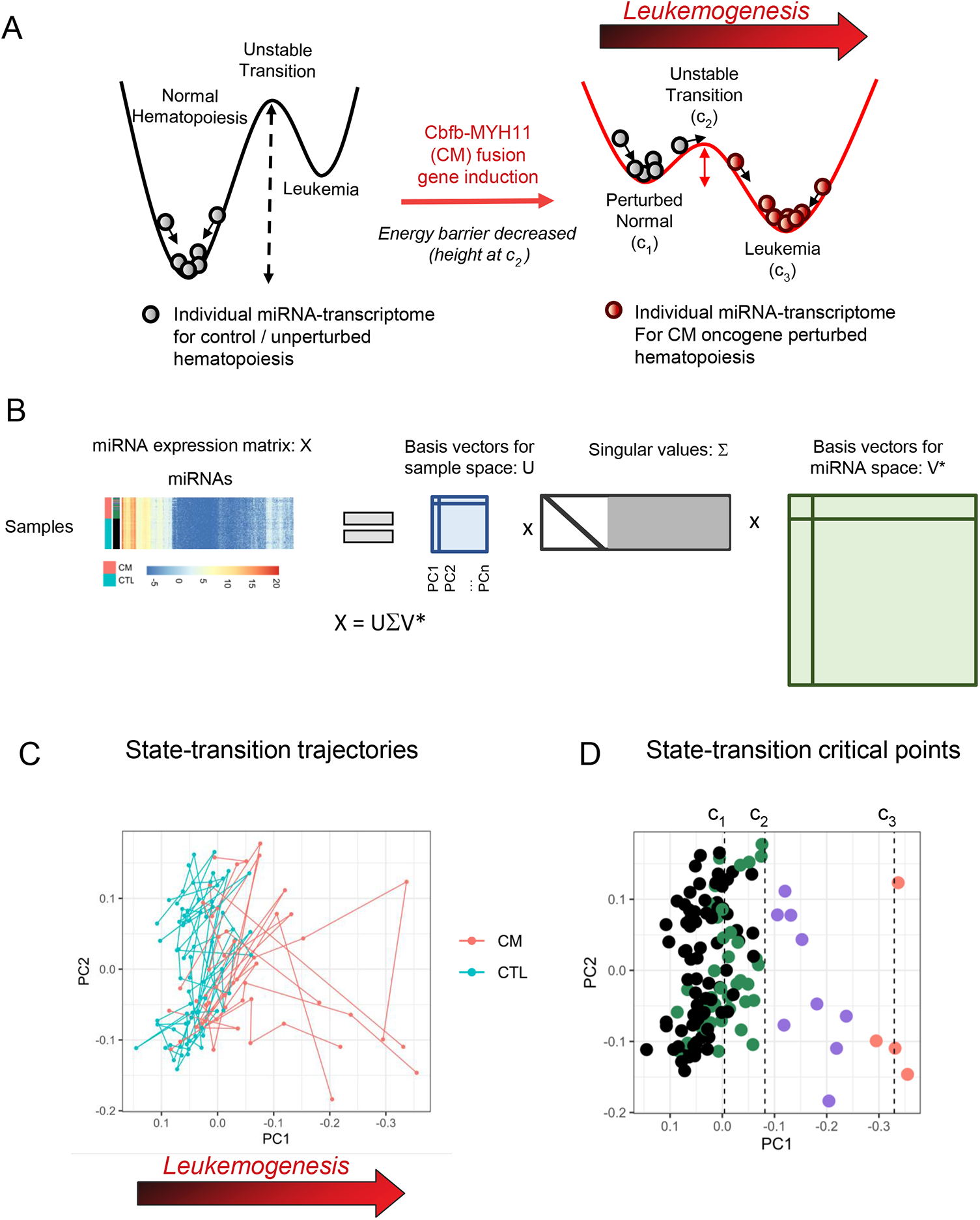
**A**. Our state-transition model represents the miRNA-transcriptome as a particle undergoing Brownian motion in a potential energy landscape. The time evolution of the miRNA transcriptome is represented as movement in the potential. In absence of an oncogenic event, there exists a large energy barrier between a state of normal hematopoiesis and leukemia, corresponding to a low probability of state-transition. An oncogenic event, such as Cbfb-MYH11 (CM) fusion resulting from inversion 16 translocation, alters the potential energy landscape, reducing the energy barrier and increasing the probability of state-transition. The states are characterized by local maxima and minima in the energy landscape, labeled *c*_1_, *c*_2_, and *c*_3_, corresponding to normal hematopoiesis, an unstable transition, and AML states, respectively. **B**. The miRNA state-space is created with the singular value decomposition (SVD) of time-sequential samples collected from control and CM induced mice over leukemia development. The SVD gives basis vectors which form principal components (*U*) representing the sample timepoints, singular values (Σ) and basis vectors for miRNA expression corresponding to the principal components (*V\**) which define *eigen-miRNA* (first column of *V\**). **C**. The first two principal components (PC1, PC2) reveal state-transition trajectories from a state of hematopoiesis to AML. **D**. State-transition critical points are identified in the state-space which characterize the state-transition from health to AML.

We modeled the miRNA state-transition trajectories for individual mice as a particle undergoing Brownian motion in the double-well quasi-potential energy with a Langevin equation of the form 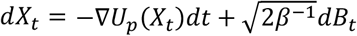. The position of the particle in the quasi-potential is denoted *X*_*t*_, *B*_*t*_ is a Brownian stochastic process that is uncorrelated in time ⟨*B*_*i*_, *B*_*j*_⟩ = *δ*_*i,j*_, and the double-well quasi-potential *U*_*p*_ is given by a quadratic polynomial specified by critical points (*c*_1_, *c*_2_, *c*_3_) which correspond to the local minima and maxima of the quasi-potential, so that *U*_*p*_(*x*) = *α*∫ (*x* − *c*_1_)(*x* − *c*_2_)(*x* − *c*_3_)*dx*, where *α* is a scaling parameter.

To be clear, we used the term quasi-potential to clarify this is a model and not a physical energy potential. We defined the double-well quasi-potential with a quadratic polynomial because this is a simple and parsimonious mathematical interpretation of our model, assuming health and AML to be stable stationary states. This assumption relies on the fact that in absence of an oncogenic event, the probability of spontaneous transition from health to AML is low, and conversely, in a state of AML, in absence of treatment, the probability of transitioning back from AML to health is also low. Although many states of the system may exist, the aim of our model was to capture state-transition dynamics between two clearly defined phenotypic states: health (i.e., normal hematopoiesis) and AML.

Because the Langevin equation of motion is a Brownian stochastic process, the probability distribution for a miRNA transcriptome particle to be at a certain position at a given time is given by a Fokker-Plank (FP) equation (**Figure S1**). Thus, to compute the probability of state-transition at a point in state-space and time *P* (*x, t*), we solved the FP equation that relates to the stochastic equation of motion, given by

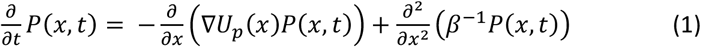

where β ^−1^ is the diffusion coefficient, and *U*_*p*_(*x*) is the quasi-potential. The diffusion coefficient was estimated for the CM and control mice as the average slope of the mean-squared displacement in the state-space over time (supplemental methods **Figure S2**).

### Constructing the leukemogenic state-space

In order to construct a miRNA-based leukemogenic state-space required to describe the miRNA transcriptome state-transition, we used the singular-value decomposition (SVD) method to perform principal component analysis (PCA) on the data matrix (X). The data matrix was composed of all time-sequential samples from CM and control mice as rows and miRNAs as columns, so that *X* = *U Σ V \** where the columns of *X* were mean-centered, log-normalized counts and * indicates the conjugate transpose. The SVD therefore decomposed the log-transformed normalized miRNA count matrix into singular values and left- and right-singular vectors (U and V* respectively; **Figure 1B**).

The singular vectors are basis vectors of the miRNA transcriptome and span either the state-space (U; composed of samples) or the feature space (V*; in this context the feature space contains the miRNA loadings and is hereafter referred to as the “miRNA space”). Therefore, singular vectors or linear combinations of singular vectors represent lower dimensional representations of the miRNA transcriptome. Additionally, each singular vector *i* has an associated singular value (σ _*i,i*_from the diagonal matrix Σ), which indicates what fraction of the total variance is explained by the associated singular vector and were ordered from largest to smallest. The associated left-singular vectors (columns of U) corresponded to the state-space and the principal components (PCs) were given as *PC* = *UΣ*. The right singular vectors (V*) were the principal component loadings and were used to define *eigen-miRNAs* (i.e., the coefficient weights of miRNA contributions to the PCs; **Table S1**).

## Results

### Construction of an AML state-transition state-space using time-sequential miRNA data and mapping the critical points of the AML double-well quasi-potential

We performed SVD on the miRNA transcriptome to construct an AML state-space where the critical points of the leukemogenic potential could be mapped and where the state-transition trajectories could be visualized. Each PC produced by SVD was correlated with the expression of *Kit* which in this mouse model, is an immunophenotypic marker of AML blasts that progressively increases during leukemogenesis. We identified PC1 as the PC that most strongly correlated with *Kit* expression (R^2^ = 0.68; p<0.001; **Figure S3; Table S2**) and that revealed the greatest separation between control and CM samples. Strikingly, as early as 1 month post induction, differences in PC1 between CM and control could be detected, prior to any evidence of leukemic cells in the peripheral blood (see **Figure S4**). Therefore, we used PC1 to define the AML state-space and AML eigen-miRNAs. To map state-transition trajectories for each mouse, we plotted time-sequential samples in a two-dimensional space constructed with PC1 vs PC2 (**Table S3**). Since each PC is orthogonal to each other by construction, any other PC would create an orthogonal 2-dimensional space; PC2 was chosen for simplicity and convenience. In the AML state-space, as the mice developed leukemia, their PC1 coordinate decreased (**Figure 1C**). Critical points (*c*_1_, *c*_2_ and *c*_3_) and AML state-transition dynamics in the state-space were identified using PC1 (see **Methods** section) and used to define states of health, transition, and AML (**Figure 1D**).

To identify the three critical points of the double-well quasi-potential *U*_*p*_ in the AML state-space, k-means clustering with k=3 was performed on the sample coordinates of PC1. The health and AML states corresponding to *c*_1_ and *c*_3_ (**Figure 1D**) were taken as the means of the clusters including the health and AML samples, denoted K1 and K3, respectively^4^. The transition critical point *c*_2_ was estimated by maximizing the Boltzmann ratio between the *c*_1_ and *c*_3_ states as 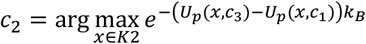 where K2 is the cluster between K1 and K3, and _*B*_ is the Boltzmann constant (**Figure S5; Table S4**). Simulation studies confirmed that the cluster means of K1 and K3 were the best estimators of *c*_1_ and *c*_3_ respectively, and *c*_2_ was best estimated to be near the boundary of clusters K1 and K2 (**Figure S6**).

### State-transition critical points enable interrogation of differentially expressed miRNA during AML development

Although the time-series sampling allowed us to observe changes in miRNA expression during leukemogenesis, the mice were not synchronized and did not develop leukemia at the same time (**Figure 2A**). Consequently, the same time points did not coincide with identical phenotypic states of AML development and therefore, could not be used to compare miRNA expression changes. Thus, to identify differentially expressed miRNAs (DE miRNAs) that occurred at the same states of AML development, we used the critical points of the double well quasi-potential identified in the AML state-space to align sequential homogeneous and therefore comparable phenotypic states of disease among individual mice (i.e., pseudo-time points). Pairwise comparisons of miRNA expression at each critical point between the CM-induced mice and the control mice allowed us to identify early, transition, late, and persistent DE miRNA events. Early events were defined as the unique DE miRNAs that occurred post-CM induction at *c*_1_, transition events were those that occurred at *c*_2_, late events were those that occurred at *c*_3_, while persistent events were the miRNAs detected as DE at all three critical points, (*c*_1_, *c*_2_, and *c*_3_; **Table S5**; **Figure 2B**).

**Figure 2.**
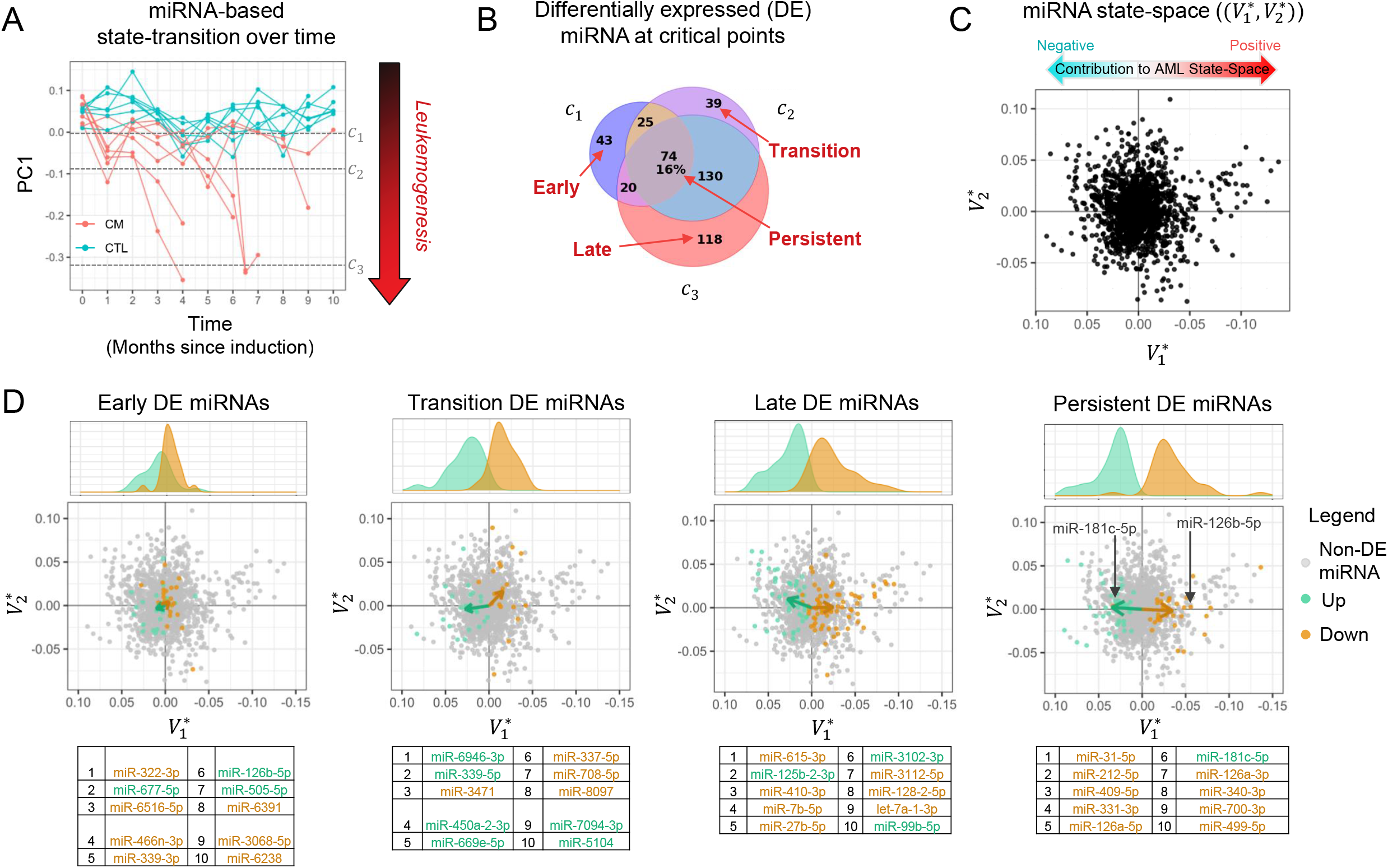
**A**. The miRNA transcriptome undergoes a state-transition from health to AML and can be seen in the first principal component (PC1) plotted over time for control and CM induced mice. All samples have a similar initial value of PC1 at time t=0 prior to CM induction. CM mice and control mice diverge and create state-transition trajectories as CM mice samples move south toward the AML state (*c*_3_). **B**. Differential expression at each critical point is used to define early, transition, late and persistent differential expression events as compared to control samples. Numbers indicate how many miRNA are differentially expressed. **C**. All miRNA are plotted in the miRNA space (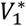 loading vs 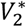 loading values) to illustrate their contribution to the AML state-space construction. The basis vectors for eigen-miRNA expression given by 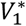 in the SVD provides a geometric interpretation of miRNA expression so that miRNA may be identified as having a positive or a negative contribution to the construction of the AML state-space. **D**. Early, transition, late, and persistent DE miRNA shown in the state-space and colored for up or down regulation. After taking the direction of expression change into account, the total contribution of the differentially expressed (DE) miRNA are visualized as a mean vector. The magnitude of the mean vector in PC1 (x-axis) indicates how strongly the DE miRNA contribute to AML state-transition (i.e., sample movement toward AML in the state-space). Early events have smaller contribution to AML, revealed by small PC1 component of the arrows, followed by increasing contributions to AML in transition events, with stronger more prominent pro-leukemogenic events indicated by larger arrows in late and persistent events. Kernel density plots above each plot also show the distribution of up- and down-regulated DE miRNA. Tables underneath each plot indicate the 10 most significantly DE miRNA of each comparison. The color of the miRNA indicates whether it was detected to be up or down regulated. The rank of each miRNA, based on their adjusted p-value, is shown in the table and used to indicate their location in the miRNA space.

Since the AML state-space is built with the eigen-miRNA loadings 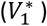, to assess the contribution of individual DE miRNAs to the leukemogenic state-space, we plotted the miRNA space (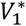 loadings vs 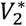 loadings; **Figure 2C**). In this newly defined miRNA space, the contribution of each miRNA to AML state-space was determined by its 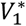 loading value (see **Figure 1B**). Thus, each miRNA had either a positive or negative contribution to the AML state-space construction. For example, miR-409-5p had the largest negative loading value, and therefore, also had the largest positive contribution to the state-space since progression to AML in PC1 went toward a more negative value (*c*_3_ AML state coordinate is negative). Whereas miR-135a-1-3p had the largest positive loading value and therefore, the largest negative contribution to the state-space.

Using the early, late, transition and persistent events, we then analyzed the contribution of the DE miRNA to movement in the state-space corresponding to AML state-transition. The contribution of each miRNA to AML state-transition depended on both its location in the miRNA space and on whether it was up- or down-regulated. A miRNA that contributed to the AML state-transition could have either: 1) a negative 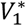 loading value and increased expression; or 2) a positive 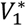 loading value and a decrease in expression. Using two miRNAs from the late events as an example, miR-126a-3p had a negative 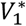 loading value (right of x=0) and was up-regulated (orange), indicating that it contributed to the AML state-transition; alternatively, miR-181c-5p had a positive 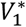 loading value (left of x=0) and was down-regulated (green), indicating that it also contributed to the AML state-transition (**Figure 2D**).

To summarize the net leukemic contribution of early, transition, late, and persistent events, we plotted vectors representing the mean of the miRNA state-space coordinates (i.e., 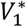 loading vs 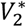 loading values) separately for the up- and down-regulated DE miRNA (**Figure 2D**). The DE miRNAs in each comparison more strongly contributed to leukemogenesis if the representative vector had a large 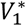 component. With this interpretation, we observed that the early events have a smaller net contribution to leukemogenesis as compared to the transition or late events. To interrogate the biological role of the DE miRNAs in leukemogenesis, we also identified pathways implicated by the DE miRNAs of each comparison^10^. Early events were involved in IL-7 signaling and metabolic pathways; transition events in Wnt and inflammation pathways; late event in MAPK and adhesion pathways, and persistent events in apoptosis, cytokines and PI3K-AKT signaling pathways (**Figure S7**).

### Dynamics of miRNA expression

To investigate changes of the miRNA patterns of expression during AML state-transition, we leveraged both the time-series samples and the critical points. Correlation coefficients between each miRNA were calculated to produce a correlation matrix. Hierarchical clustering of the correlation coefficients revealed four distinct groups of miRNA expression dynamics (**Figure 3A; Table S6**). When plotted in the state-space (PC1), the miRNA expression dynamics corresponded to two patterns: non-monotonic (**Figure 3B**; groups 1,3); and monotonic (**Figure 3B**; groups 2,4). The miRNA groups which exhibited a non-monotonic expression dynamic revealed a local maximum (group 1) or local minimum (group 3) around the transition critical point *c*_2_. As *c*_2_ is the unstable stationary point in the state transition model that separated healthy and AML states, changes in expression at *c*_2_ indicated that these miRNAs likely play a role in facilitating the irreversible transition from health to AML. Without the identified critical points to align the leukemogenic states of time-series samples, the nonlinear expression patterns of group 1 and 3 would not be detected or interpreted in this way.

**Figure 3.**
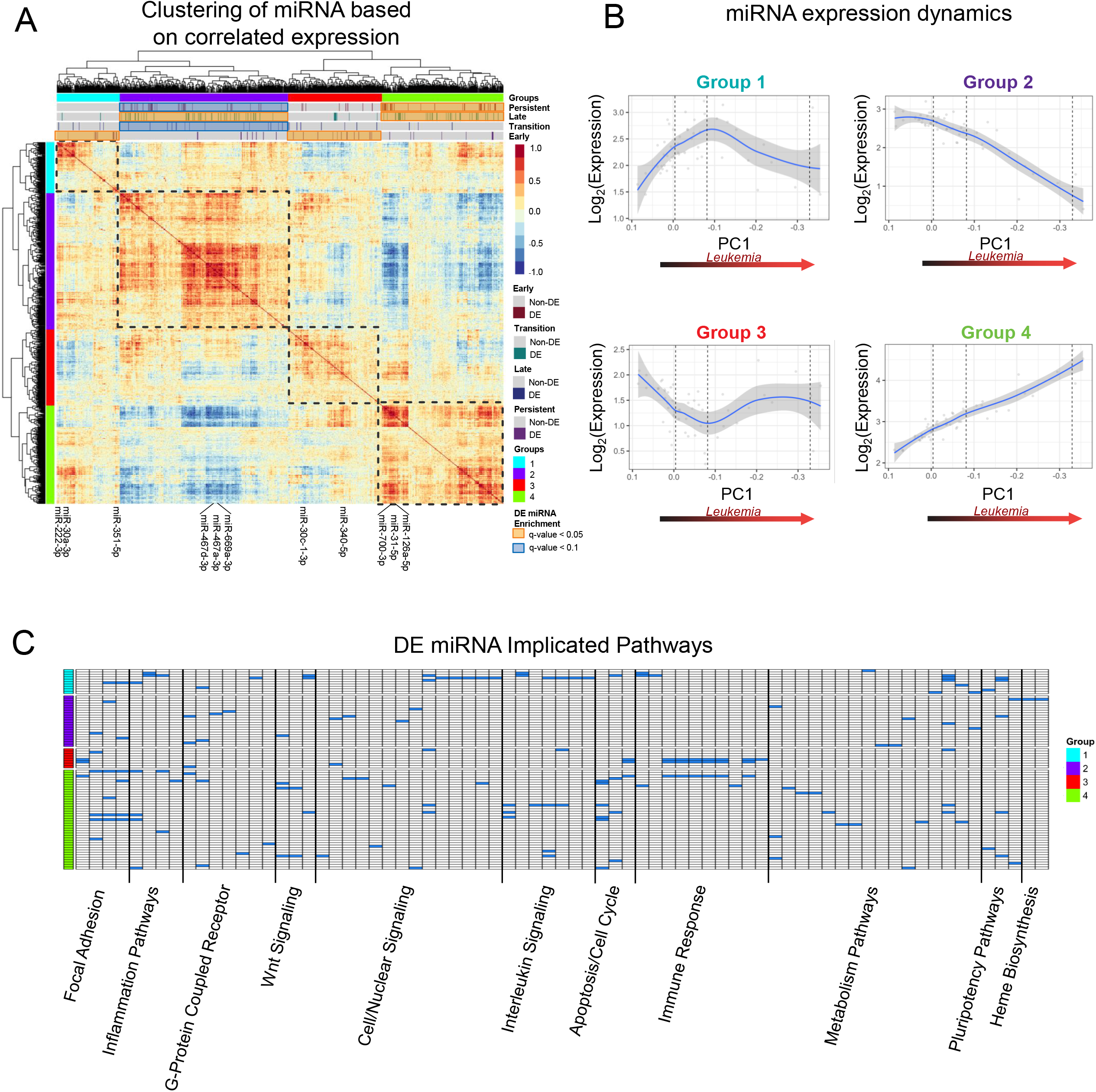
**A**. Hierarchical clustering of correlated miRNA expression annotated with state-transition critical point DE miRNA. Four distinct patterns of miRNA expression are identified. **B**. Four patterns, or groups of miRNA expression dynamics include monotonic (groups 2,4) and non-monotonic (groups 1,3) patterns when plotted in the state-space (PC1). Interestingly, a local maximum (group 1) and minimum (group 3) are identified very near the unstable transition critical point *c*_2_. **C**. Pathways implicated based on miRNA expression dynamics are summarized to reveal the non-monotonic pattern in group 1 is associated with both Cell/Nuclear Signaling pathways including IL-6, TNFα-NFκB signaling, and cytokines inflammation response, and Wnt signaling. The opposite non-monotonic expression dynamic (group 3) is enriched for Immune Response pathways including antigen processing and toll-like receptor signaling. Full list of pathways in each summarized category is shown in **Table S7**.

Of note, when we tested each group for over-representation of DE miRNAs using the hypergeometric test, we observed that the monotonic groups 2 and 4 showed over-representation of late and persistent events respectively whereas the non-monotonic groups 1 and 3 showed over-representation of early events. Pathways implicated based on miRNA expression dynamics reveal the non-monotonic pattern in group 1 was seemingly associated with IL-6, TNFα-NFκB signaling, cytokines inflammation response, and Wnt signaling while group 3 was seemingly associated with immunologic pathways including antigen processing and toll-like receptor (TLR) signaling (**Figure 3C;** pathway summary in **Table S7**). Similar to group 3, group 4 also was associated with immunogenic and TLR signaling but differed from group 3 in its association with PI3K-AKT signaling. Group 2 showed heterogeneous pathway involvement.

### Validation of miRNA state-space in an independent cohort

As a validation of the SVD derived state-space, we used an independent cohort of CM and control mice (validation cohort). The eigen-miRNA loadings 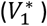 were used to project data from the validation cohort into the state-space. Without any prior knowledge of the genotype or timepoint of the samples, the eigen-miRNAs predicted the disease status and trajectories of the validation samples (**Figure 4A**). Of note, the state-space trajectories of three CM mice in the validation cohort which were induced but did not develop AML during the observation period (6 months) were correctly projected to be with the control mice (black arrows; **Figure 4A**).

**Figure 4.**
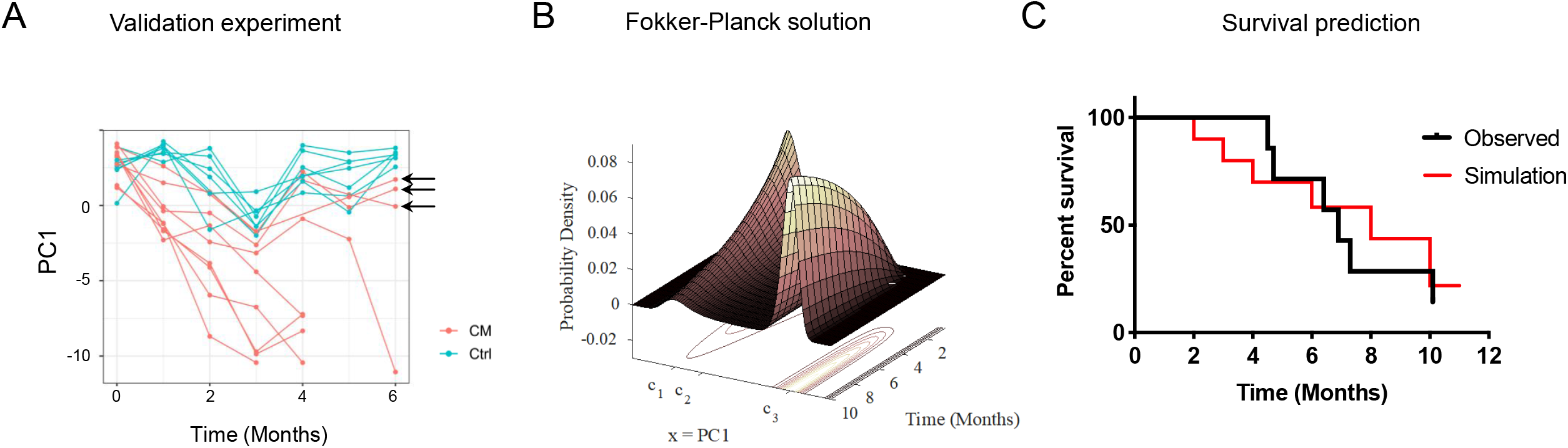
**A**. State-space predictions on an independent validation experiment. Using only the coefficient weights determined by the eigen-miRNA in V*, new samples are projected into the state-space to characterize state-transition trajectories. Control and CM trajectories are accurately identified, including 3 CM induced mice which did not develop leukemia at the experiment endpoint (t=6) indicated with arrows. **B**. The solution of the Fokker-Planck model of state-transition gives the evolution probability density function (PDF) and prediction of state-transition for any point in the state-space at any time. **C**. The PDF is used to predict the time to develop AML from an initial state of the miRNA transcriptome and accurately predicts the manifestation of leukemia in the mice.

### The state-transition model correctly predicts time to AML development

Thus far we have illustrated the use of pseudo-time points (i.e., critical points) to phenotypically synchronize the transition of each mouse from health to AML. However, the state-space model can also be used to predict the AML development in each mouse in real time. To this end, in order to predict the time to AML, we initialized the FP equation (**Eq. 1**) with a Gaussian distribution based on the first time point post induction (t=1) of CM samples in the state-space and solved the equation forward in time (**Figure S1**). By integrating the solution of the FP from *c*_2_ to *c*_3_, we calculated the probability of state-transition over time. We then compared the predicted time to develop AML with that observed in the mice using the Kaplan-Meir estimator. Our model accurately predicted the time to develop leukemia for the cohort of mice, as the predicted and observed survival curves were not significantly different (p>0.05; **Figure 4C**).

### mRNA and miRNA transcriptomes both encode a similar but not identical AML state-transition

We show here that the miRNA transcriptome undergoes state-transition during leukemogenesis similar to the mRNA transcriptome (RNA-seq) that we previously reported using the same mouse model^4^. However, in contrast to the miRNA-derived state-space, which was encoded in PC1, mRNA state-transition was encoded in PC2. Nevertheless, the overall displacement of each mouse in the state-space, defined to be a connecting line between the first (t=0) and the last time points, revealed similar trajectories in both the mRNA and miRNA AML state-spaces (**Figure 5A; Table S8**).

**Figure 5.**
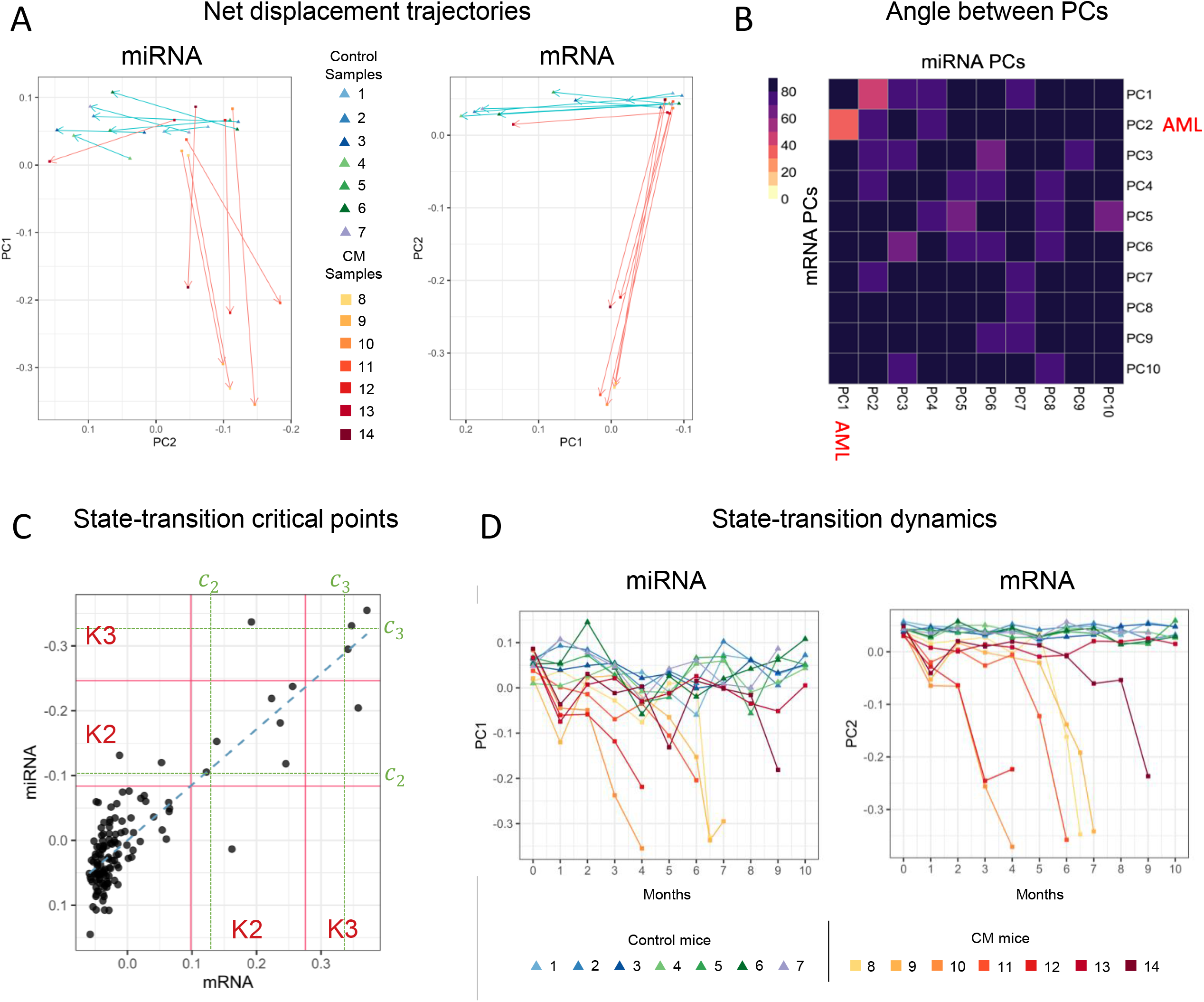
**A**. By plotting the first and last timepoint samples in miRNA and mRNA state-spaces, net displacement trajectories reveal similarities between the gene expression and miRNA state-transition state-spaces. **B**. The angle between mRNA (x-axis) and miRNA was computed for every pair of principal components. The smallest angle, which indicates the states with the most similarity, between all states was detected for the AML state-space (miRNA PC1 and mRNA PC2). The close alignment of the miRNA- and mRNA-derived AML state-spaces indicate that the time-evolution of AML state-transition is more similar than any other source of variance. **C**. State-spaces for miRNA and mRNA plotted against each other annotated with critical points reveals similarities in both the state-transition dynamics and the critical point classification. Only 5 out of 132 samples would be categorized differently depending on which critical points were used, mRNA or miRNA. **D**. The trajectories for both the miRNA and mRNA AML state-spaces for each sample are plotted as function of time.

To quantitatively assess the similarity and differences of the mRNA- and miRNA-derived state-spaces, we then computed the angle between all the miRNA and mRNA transcriptome PCs pairs using the vector dot product (see supplemental methods). We interpreted the results as follows: miRNA and mRNA transcriptome PCs that were orthogonal to each other encoded different sources of variation within each data type; whereas miRNA and mRNA transcriptome PCs that had an angle closer to zero encoded sources of variance that were shared between the two data sets. Strikingly, these two PCs with the smallest angle, and therefore the most similar, were those that encoded the variation associated with state-transition from health to AML (i.e., PC1 for miRNA and PC2 for mRNA; **Figure 5B**). Not only were no other PCs more aligned than miRNA PC1 and mRNA PC2, but the other PCs were also nearly orthogonal to each other (**Figure S8**). This suggests that the time-series dynamics of mRNA and miRNA can both be used to predict AML development since they encode similar, albeit not identical, information of the state-transition from health to disease. In fact, by plotting the miRNA and mRNA state-spaces against each other and annotating the critical points, the overall state-transition trajectories had a striking similarity (**Figure 5C,D**). Only 5 samples out of 129 total samples were classified differently as being in *c*_1_, *c*_2_, or *c*_3_ states based on which critical points—mRNA or miRNA—are used.

Taken together, state-transition trajectories, the angle between the principal components, and the locations of the critical points between mRNA and miRNA-derived state-spaces, we concluded that both miRNA and mRNA expression undergo a system-wide state-transition during AML development and they encode information of disease progression that is similar, but not identical for the leukemogenic processes.

## Discussion

Here we report the application of a state-transition theory to the interpretation of how temporal changes in the miRNA expression informs AML initiation and progression. Using miRNA expression analysis of PBMCs collected at sequential time points from a mouse model of inv(16) AML from induction of the leukemogenic CM fusion gene until development of overt disease, we identify “key” states of the miRNA transcriptome corresponding to the critical points of a miRNA-based leukemic double-well quasi-potential. To confirm the accuracy of our prediction, we utilized time-series miRNA sequencing data from an independent cohort of mice. Notably, we found no difference when we compared the predicted vs actual survival curves, supporting the accuracy of our state-transition model prediction.

The state-transition model is a system-wide holistic approach to biology where the transformation from one state to another is viewed as a change in state of the whole system as opposed to a change resulting from one (or a small collection) of molecules^5,11,12^. Using gene expression as an example of this perspective, a state transition occurs because the entire mRNA transcriptome transitions to a new steady state: the expression of all genes contribute to the transition, not the change in expression of a single gene. To be clear, as was the case in our CM AML model, a single mutation or molecule may be sufficient to cause a perturbation that induces a state-transition; however, in the system-wide holistic view, the perturbation causes an alteration to the underlying gene regulatory network that results in a state-transition to the entire transcriptional state. Currently, miRNA expression is more commonly used to investigate the regulatory effects of a single or small set of miRNA molecules. Although smaller in dimensionality, the miRNA transcriptome, similar to the mRNA transcriptome, is both involved in a wide range of biological processes and a highly regulated subset of RNA molecules. Our finding that the miRNA transcriptome can be used to construct an AML state-transition model is, to the best of our knowledge, the first report of miRNA transcriptome encoding system-wide dynamics during the course of AML pathogenesis and progression. Thus, this work reveals that a more system-wide holistic view of the miRNA transcriptome is warranted.

The state-space and state-transition model provided a theory-guided approach to the analysis of differential expression, with early, transition, late, and persistent events defined relative to critical points in the state-transition. The biological significance of the critical points is provided by the analysis of early, transition, late and persistently DE miRNAs that are respectively associated with *c*_1_, *c*_2_, *c*_3_ or all three critical points. This in turn allowed for a novel approach to quantify miRNA contributions to AML pathogenesis through identification of distinct dynamics of miRNA expression, including monotonic and non-monotonic patterns. The miRNAs groups (2 and 4) with a monotonic patterns of expression changes, i.e., continuously decreased expression (group 2) or continuously increased expression (group 4) were enriched with miRNAs that were found to be persistent DE miRNAs at all three critical points. These groups included miRNAs that regulate genes involved in the “inflammasome” (i.e., miR-467), cell differentiation (i.e., miR-669, miR-31) and leukemia stem cell function (i.e., miR-126). The miRNA groups with a non-monotonic expression patterns included several miRNAs that regulate glucose and lipid metabolisms (i.e., miR-320 and miR-142) or directly target KIT (i.e., miR-122) or ubiquitination (i.e., miR-378, miR-30c). While the biological meaning of the two patterns of expression dynamics requires additional analyses and experimental evidence both *in silico* and at the bench, it is possible that they may represent key features of leukemogenesis. We interpreted the monotonic patterns of expression dynamics as the representation of a “leukemogenic force” given by the continuous increase and decrease of onco- and tumor-suppressor miRNAs respectively during AML state-transition. In contrast, it is possible that the non-monotonic patterns of expression dynamics represent a “restoring force” that attempts to return the miRNA-transcriptome to the initial equilibrium (i.e., health or *c*_1_) after CM induction but that inevitably breaks down at the point of “no-return” in AML state-transition (i.e., the critical point *c*_2_), where it is overcome by the “leukemogenic force”. This work therefore suggests that the dynamics of miRNA transcriptome encode critical information and may serve as a novel blood-based biomarker of state-transition from health to AML or an early indicator for response to treatments (i.e., transition from AML to health) upon a therapeutic intervention.

Of note, we recently showed that state-transition theory and double-well potential of the mRNA transcriptome can also predict AML initiation and development.^4^ When we analyzed state-transition trajectories, the angle between the principal components, and the locations of the critical points between mRNA and miRNA-derived state-spaces, we observed that both miRNA and mRNA expression undergo a system-wide state-transition during AML development and they encode information of disease progression that is similar, but not identical. Thus, while miRNAs and mRNA are mostly functionally associated, it is possible that certain steps of leukemogenesis are instead uniquely dependent on either miRNA or mRNA expression. We expect that simultaneous state-transition modeling of miRNA and mRNA expression dynamics will provide a unique perspective to map the inter-relationships and information content that will be instrumental to detect early indications, monitor treatment response, and predict relapse in individual AML patients.

## Acknowledgements

This work was supported in part by the NIH under award number R01CA178387 (to Y.-H. Kuo), R01CA205247 (to Y.-H. Kuo/G. Marcucci), U01CA25004467 (to R.C. Rockne, Y-H. Kuo, G. Marcucci), R01CA248475 (G. Marcucci), T32CA221709 (D.E. Frankhouser), and the Gehr Family Center for Leukemia Research. Research reported in this publication included work performed in the Integrated Genomics Core, Bioinformatics Core, Analytical Cytometry Core, and Animal Resource Center supported by the NCI of the NIH under award number P30CA33572. The content is solely the responsibility of the authors and does not necessarily represent the official views of the NIH.

## Supplemental materials and methods

### miRNA-seq library preparation and sequencing

Samples were allocated to randomized batches for library preparation, such that samples from each timepoint were distributed evenly over all sequencing runs. All libraries were prepared using the Illumina TruSeq Small RNA protocol with minor modification following the manufacturer’s instructions. Briefly, for each sample, 280 ng of total RNA was ligated to the sRNA 3′ adaptor (5’-TCTGGAATTCTCGGGTGCCAAGGAACTCC-3’) with T4 RNA Ligase 2, truncated (New England BioLabs) for 1 h at 22°C, and subsequently ligated to a 5′ adaptor: 5’-GUUCAGAGUUCUACAGUCCGACGAUCNNN-3’) with T4 RNA ligase 1 (New England BioLabs) for 1 h at 20°C. The constructed small RNA library was first reverse-transcribed using GX1 (5′-GGAGTTCCTTGGCACCCGAGA) as the RT primer then subjected to PCR amplification for 13 cycles, using the primers GX1 (5′-CAAGCAGAAGACGGCATACGAGAT[NNNNNN]GTGACTGGAGTTCCTTGGCACCCGAGAATTCCA-3’) and GX2 (5′-AATGATACGGCGACCACCGAGATCTACAC[NNNNNNNN]CGACAGGTTCAGAGTTCTACAGTCCGA-3’), followed by 6% TBE PAGE gel purification with size selection (for targeted small RNAs of 17–35 nt). Individual libraries were prepared using a unique index primer (NNNNNNNN in the GX1 and GX2 primer) in order to allow for pooling of multiple samples prior to sequencing. The purified libraries were quantified using qPCR. Sequencing of single end 50 cycles was performed on a HiSeq 2500 (Illumina Inc., San Diego, CA), and image processing and base calling were conducted using Illumina’s RTA pipeline.

Raw sequencing reads were processed with the nf-core smRNASeq pipeline version 1.0^13^ using the GRCm38 genome reference (with the parameter --genome GRCm38) and adapters for the Illumina small RNA protocol (by setting --protocol illumina). Briefly, trimmed reads were mapped using bowtie^14^ to miRBase^15^ mature miRNAs (using the parameters -k 50 --best --strata), and the number of reads mapping to each was counted using samtools stats^16^. Each library was also subjected to extensive quality control, including estimation of library complexity, contamination, sequence quality, read length and depth, among other metrics detailed in the pipeline repository. Mapped reads were merged into a matrix of counts per gene for each sample at each timepoint and normalized to counts per million (CPM) reads mapped, as implemented in edgeR^17^. Surrogate variable analysis was used to check for confounding experimental effects^18^. None were apparent (data not shown). The miRNA dataset is submitted to GEO and accession number pending.

### miRNA analysis

Log normalized miRNA were generated by taking the log (base 2) of CPMs [i.e., log2(CPM+0.01)] and used for singular value decomposition (SVD). We treated each mouse as a replicate to investigate how expression changed as the mice moved through the leukemic state space (PC1). Differentially expressed (DE) miRNA were determined by comparing control samples to the sample classified as each of the critical points (*c*_1_, *c*_2_, and *c*_3_) using miRTOP generated miRNA counts and default settings of DEseq2^19,20^ (**Figure 2B-D**). For the validation cohort, data were processed by removing adapters using cutadapt v1.9.1, and trimmed sequences were aligned to mm9 genome using Bowtie v0.12.7 with “--best” option^21,22^. Mature miRNAs counts were determined using R scripts and miRbase v21^15^. Log normalized counts were again generated from CPM [i.e., log2(CPM+0.01)]; one sample (out of 99 total samples) was removed as an outlier based on poor library quality and abnormal expression patterns. To project the validation cohort samples into the AML state-space, the log normalized expression matrix (*X*_*V*_) was multiplied by (i.e., *U*_*V*_ = *X*_*V*_ * *V*), and time was plotted vs the first component of *U*_*V*_ (**Figure 4A**).

### Identification of leukemia state-transition state-space and comparison to *Kit* expression

In order to identify which principal component was most associated with leukemia, we examined all principal components and correlated them with expression of *Kit* gene, which is an immunophenotypic marker of AML. PC1 had both the highest R^2^ and lowest p-value of all PCs (**Table S2**; **Figure S2**). PC1 and PC2 accounted for 5 and 4%, respectively, of the total variance present in the data. *Kit* expression was determined using the matched mRNA sequencing (RNA-seq) for each sample which is previously described^4^.

### Angle between mRNA and miRNA principal components

The angle between each of the PCs from mRNA and miRNA (**Figure 5B**) is computed such that for two vectors **a** and **b**, 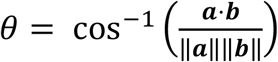 where ‖⋅‖ is the L-2 norm, or magnitude of the vector. The angle is computed using all PCs with non-zero eigenvalues for the full state space with all samples.

## Notes

### Competing Interest Statement

The authors have declared no competing interest.

