## Supplemental Material for "State-Transition Analysis of Time-Sequential microRNA Expression Predicts Development of Acute Myeloid Leukemia"

Ya-Huei Kuo, Russell C. Rockne, Guido Marcucci

1500 E. Duarte Rd.

Duarte, CA 91010

**Supplemental Tables;** *- denotes separate file

1. Eigen-miRNA*
2. State-space coordinates*
3. Kit expression correlation*
4. Model and parameter values
5. DE miRNA lists*
6. miRNA in each expression dynamics group*
7. Pathway summary*
8. Sample IDs*

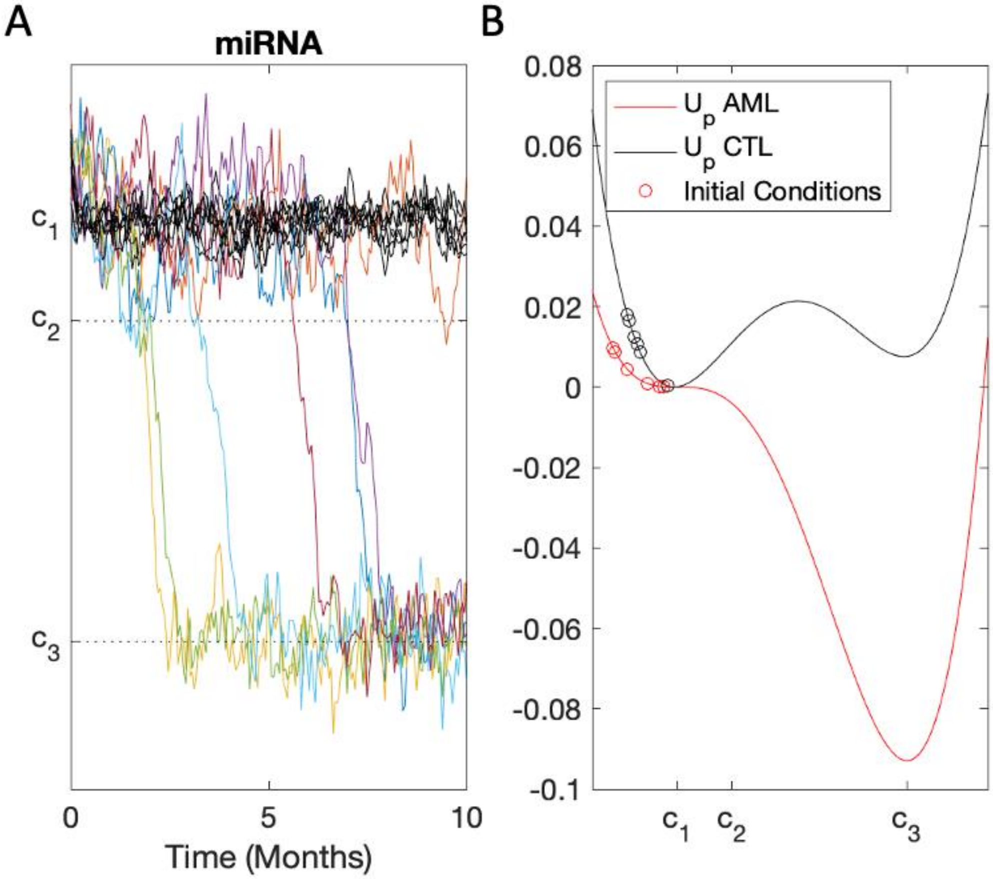


Figure S1: Simulation of miRNA state-transition dynamics. A. By solving the Langevin equation of motion forward in time $dX_t = -\nabla U_p(X_t)dt + \sqrt{2\beta^{-1}}dB_t$ with parameters and initial conditions (t=1 month post induction) estimated from the data, the model correctly predicts the transition from a state of normal hematopoiesis ($c_1$) to AML ($c_3$) with different mice (colored lines) manifesting AML at different time points. Controls (black lines) remain in the normal state of hematopoiesis ($c_1$). B. The quasi-potentials inferred from the data for CM mice undergoing state-transition to AML (red line) and controls (black line). The shape of the AML potential clearly shows a predicted state transition to the lower energy state $c_3$ as compared to the control potential.

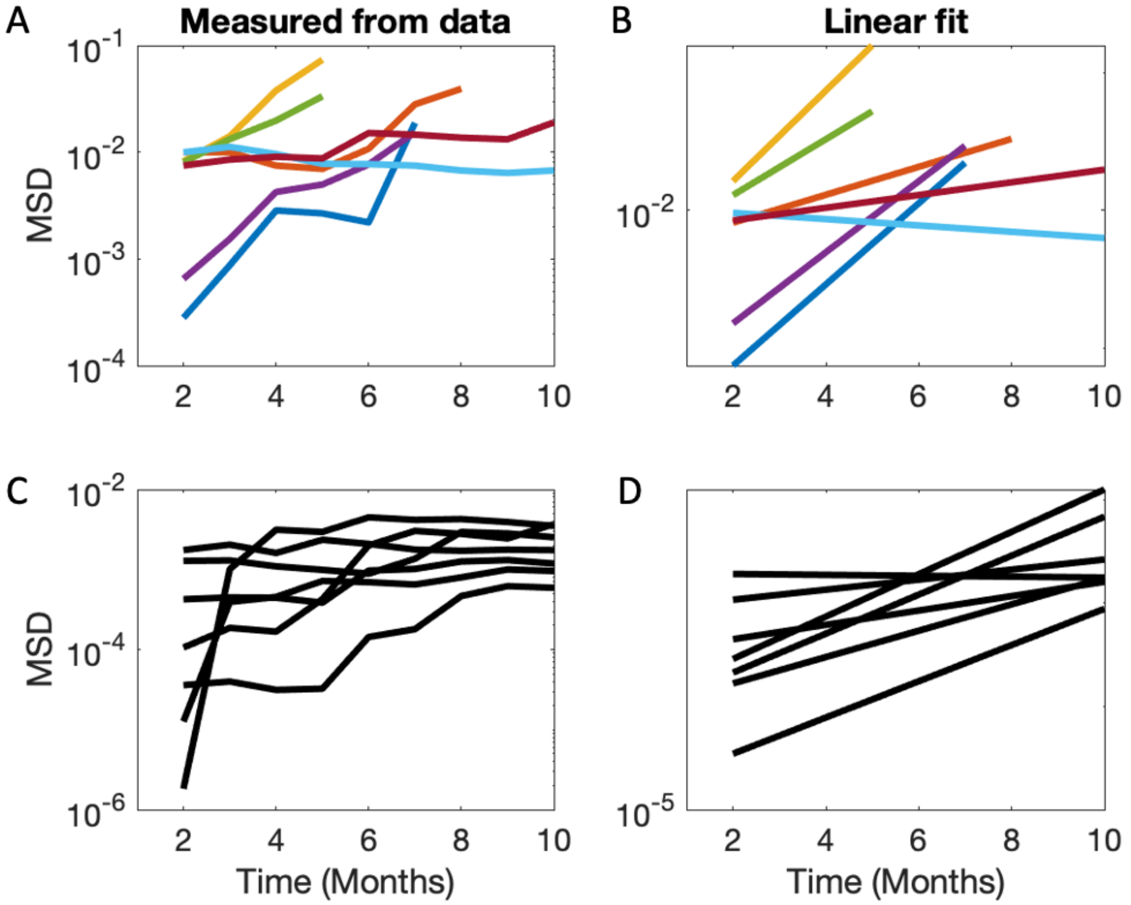


Figure S2: Mean squared displacement (MSD) analysis of miRNA trajectories in state-space. Top. MSD analysis of CM mice. A: MSD computed from PC1 trajectories for n=7 CM mice log-scale y-axis. Each mouse trajectory is a different color, coordinated with panel on the right. B: linear fit of the MSDs for each CM mouse. The mean of the slopes of the linear fits is used as an estimator of the diffusion coefficient $\beta^{-1}$ in both the Langevin equation of motion and Fokker-Planck probability density models. Bottom: same analysis for control mice (C,D). Note the slopes of the MSD curves and linear fits are smaller for the control mice as compared to the CM mice. The flat MSD curves and reduced slopes for the control mice suggesting confined diffusion, as compared to the CM mice.

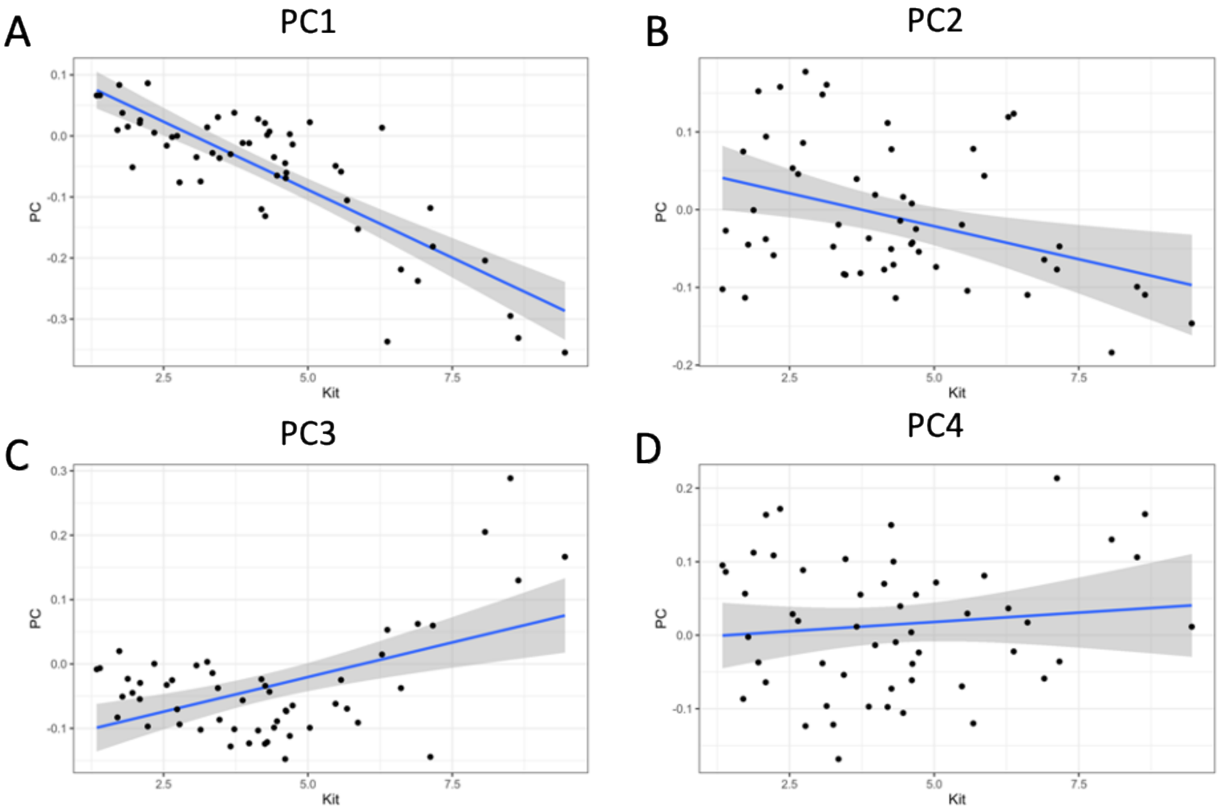


E

| PC | R-squared | p-value |
|---|---|---|
| 1 | 0.6819 | 2.75E-14 |
| 2 | 0.1432 | 0.00521 |
| 3 | 0.2518 | 0.00013 |
| 4 | 0.0125 | 0.42493 |

Figure S3: All principal components (PCs) were tested for correlation with Kit expression. A-D. The first four PCs, which were above the “elbow” of the scree plot, are shown. PC1 showed the best correlation ($R^2$=0.68; p-value < 0.001) of all PCs and was selected as the AML state-space.

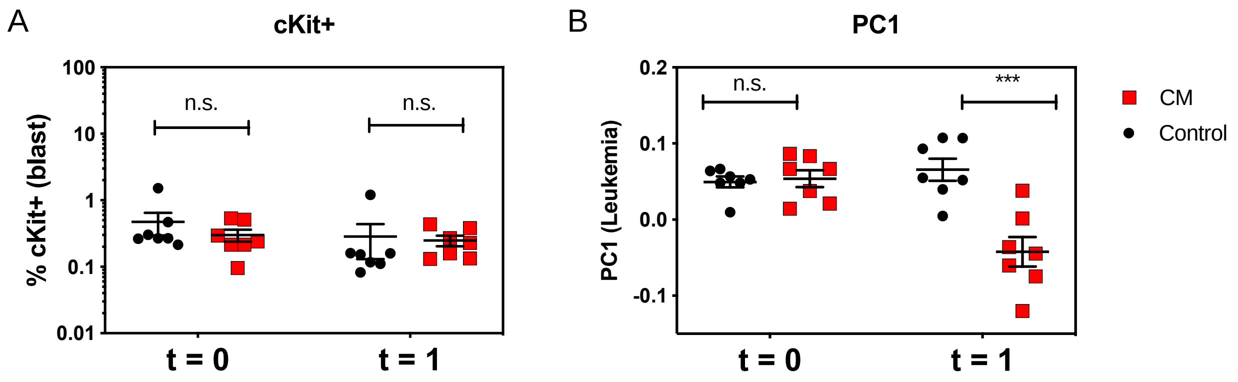


Figure S4: A. CM and control sample cKit+ cells are compared at both the pre-induction time point (t=0) and the first time point after induction of the CM fusion gene (t=1) using flow cytometry[1]. cKit+ cells are used as a marker for leukemia; no change was detected between control and CM samples at either time point. B. When the same comparison between CM and control is made using the AML state-space component (PC1), the CM samples show a significant decrease in their PC1 component post-induction. The change in PC1 component indicates that the samples have started transitioning toward AML in the state-space. Therefore, the state-space representation (i.e. the miRNA transcriptome) is an early indicator of state-transition and is able to detect transition toward leukemia before any cKit+ cells can be detected.

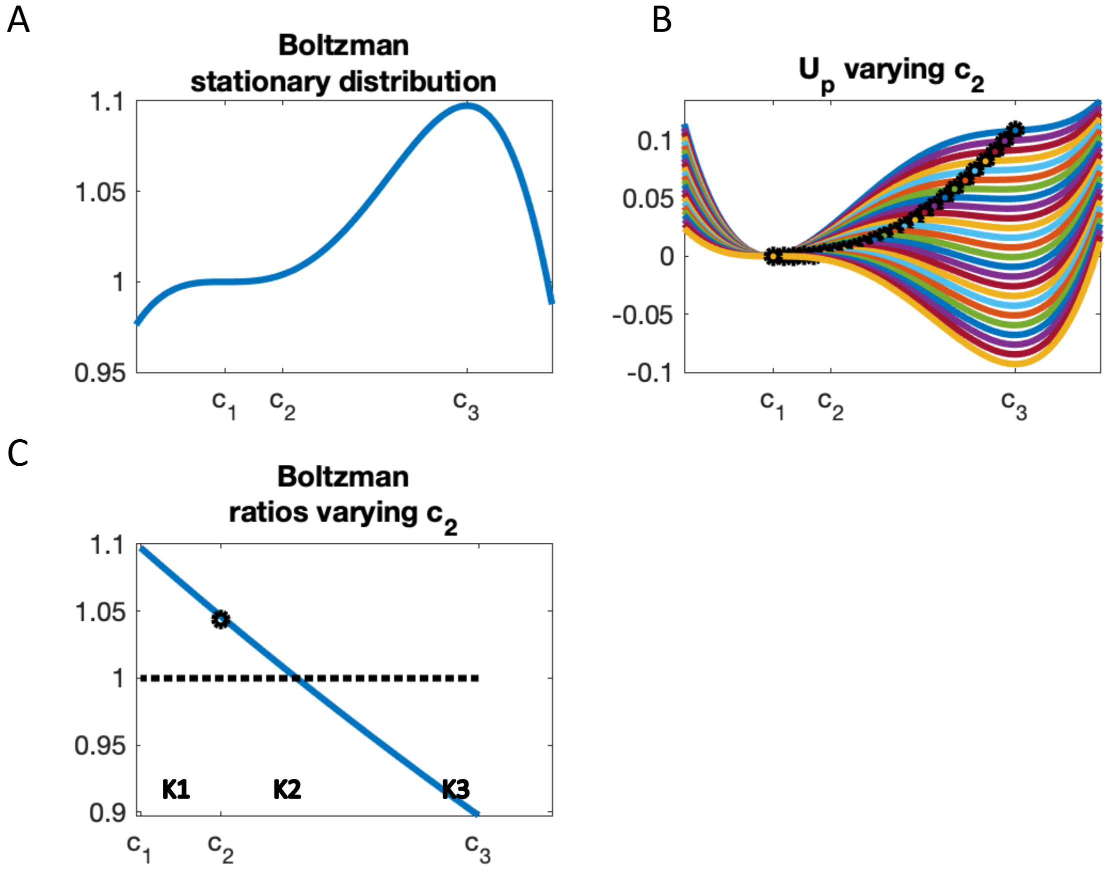


Figure S5: Estimation of the transition critical point $c_2$. A. The Boltzmann stationary distribution in the state-space is given by $p(x,\infty) = \exp\,(-\nabla U_p)$. B. The quasi-potential computed for different values of $c_2$ in the range $c_1 \leq c_2 \leq c_3$, where the location of $c_2$ is shown as a black circle. As $c_2$ varies from $c_1$ to $c_3$, the shape of the quasi-potential changes, and so too does the predicted stationary distribution. C. To summarize the probability of one state ($c_1$) or the other ($c_3$) for various values of $c_2$, we compute the Boltzmann ratio (B.R.), given by $B.R. = \frac{\exp(-\nabla U_p(c_3))}{exP(-\nabla U_p(c_1))}$. If $B.R. > 1$, then the state $c_3$ is more likely, if $B.R. < 1$ then the state $c_1$ is more likely. We see that the B.R. is maximized for the $c_3$ state at the boundary of the K1 and K2 clusters. Values of the parameters are given in Table S1.

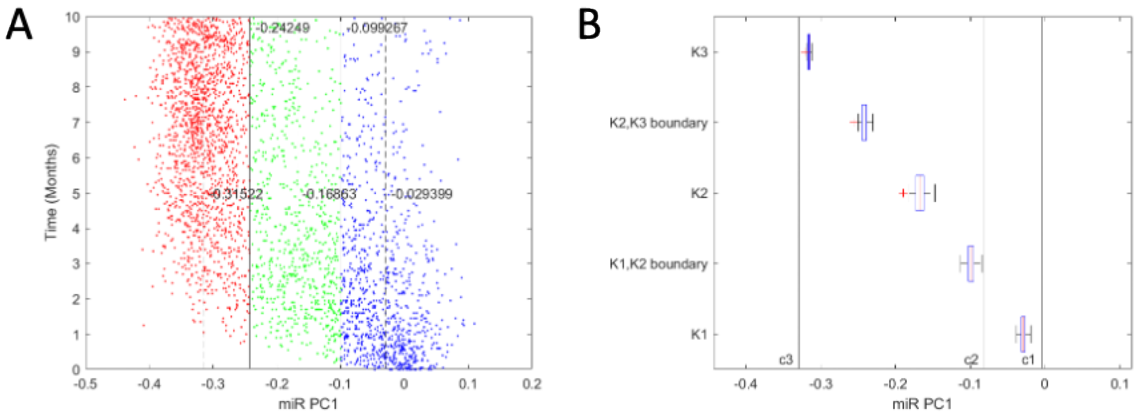


Figure S6: Simulation study of estimation of transition critical point $c_2$. A. By solving the Langevin equation forward in time to create 1000 virtual state-transition trajectories and randomly sampling 100 points from each trajectory, a virtual state-space is created over time (x-axis PC1, y-axis time). We performed k-means clustering with k=3, (K1 = blue, K2 = green, K3 = red) and plot the cluster means and boundaries between clusters K1-K2 and K2-K3. B. The critical points $c_1, c_2, c_3$ are shown as solid lines and the estimators derived from A) are shown as box and whisker plots (red line is mean, box is inner quartile, and whiskers are outer quartiles) of the distribution of samples. We see that the mean of K1 is close to $c_1$, the boundary of K1 and K2 is closest to $c_2$ and the mean of K3 is closest to $c_3$, supporting our approach to use these as estimators for the critical points in our model. Actual data is less densely sampled in the state-space and produces slightly different estimators.

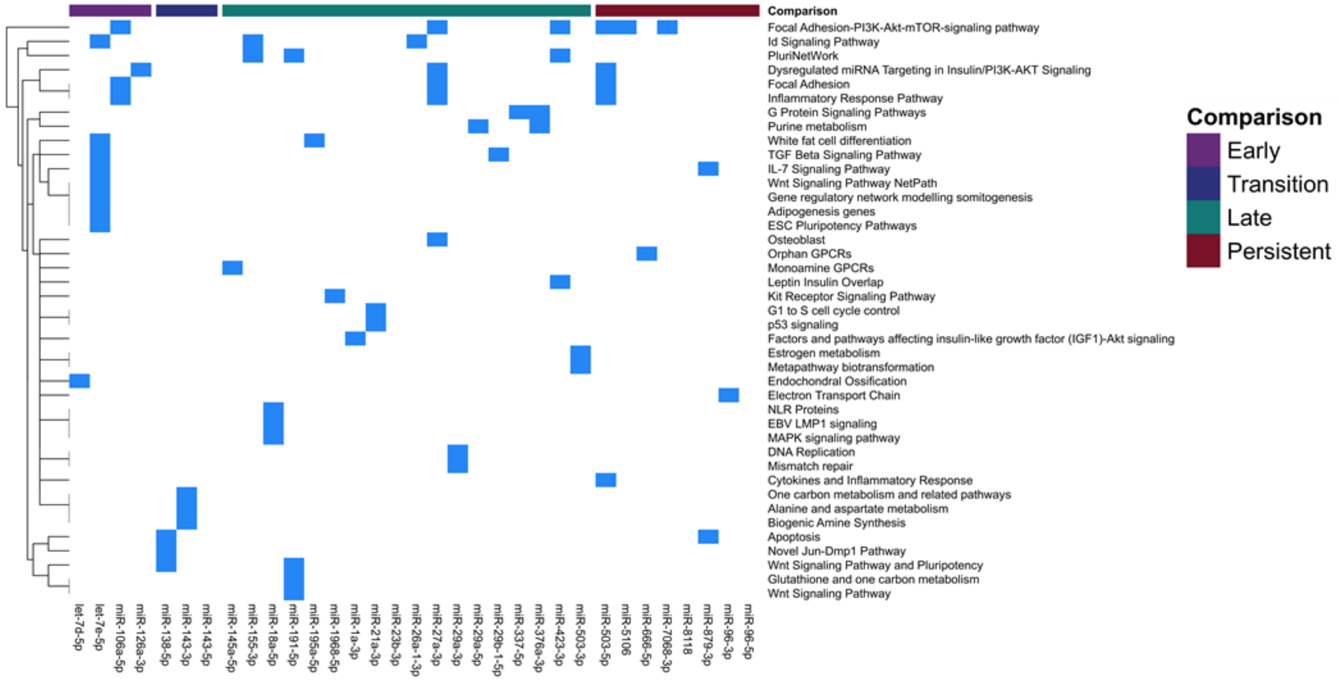


Figure S7: The pathways implicated by DE miRNA for each of the early, transition, late, and persistent events are shown by indicating the miRNA implicated in each pathway (blue). Any experimentally validated KEGG or WikiPathway pathway associated with a DE miRNA from the four comparison was reported[2].

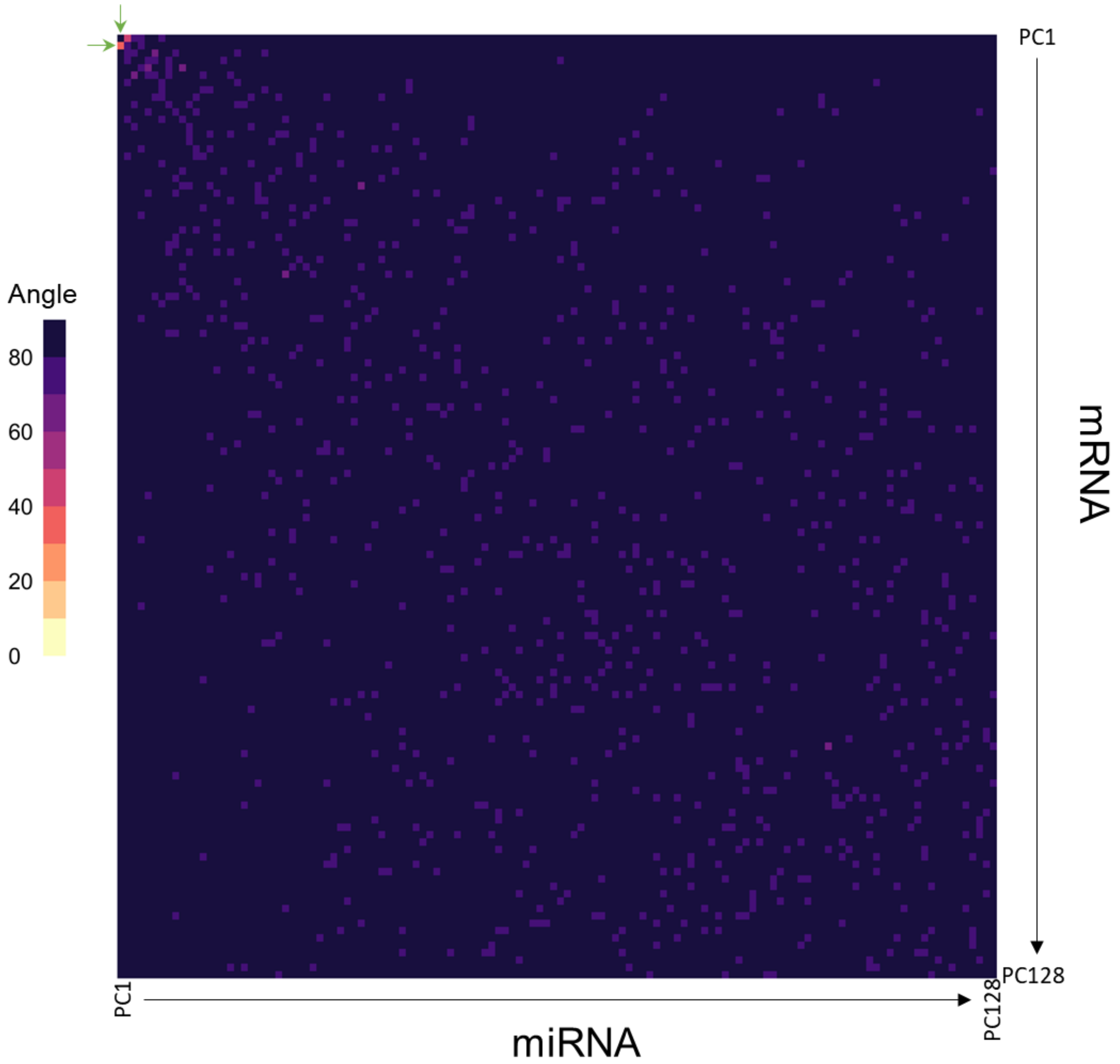


Figure S8: The angle (in degrees) is shown between all non-zero PCs from both the miRNA (x-axis) and mRNA (y-axis) data. The angle between PCs was determined by taking the inverse cosine of their dot product. An angle close to zero indicates that the two PCs from each data type represent a similar source of variation among the samples. The PCs with the lowest angle are those corresponding to the AML state-space (miRNA PC1 and mRNA PC2; green arrows). An angle close to 90$^{o}$ indicates that the two PCs represent sources of variance within the data that are orthogonal.

Table S4. Model and simulation parameters. For controls, $c_2$ is taken to be the midpoint between $c_1$ and $c_3$. Parameters estimated as described in main text and supplemental methods.

| Parameter | Meaning | Value |
|---|---|---|
| $\boldsymbol{c_1}$ | Normal hematopoiesis | -0.0042 |
| $\boldsymbol{c_2}$ | Transition state | -0.0816 |
| $\boldsymbol{c_3}$ | AML | -0.3294 |
| $\boldsymbol{\alpha}$ | Scaling factor of quasi-potential | 100 |
| $\boldsymbol{\beta}$ | Diffusion coefficient | 285.7143 |
| $\boldsymbol{\beta_H}$ | Diffusion coefficient for controls | 1.4286e+03 |
